# *Plasmodium* oocysts respond with dormancy to crowding and nutritional stress

**DOI:** 10.1101/2020.03.07.981951

**Authors:** Tibebu Habtewold, Aayushi A. Sharma, Claudia A.S. Wyer, Ellen K.G. Masters, Nikolai Windbichler, George K. Christophides

## Abstract

Malaria parasites develop and grow as oocysts in the mosquito for several days before being able to infect another human. During this time, mosquitoes take regular bloodmeals to replenish their nutrient and energy reserves needed for flight and reproduction. We hypothesized that supplemental bloodmeals are critical for oocyst growth and that experimental infection protocols, typically involving a single bloodmeal, cause nutritional stress to developing oocysts. Therefore, enumerating oocysts independently of their growth and differentiation state may lead to erroneous conclusions regarding the efficacy of malaria transmission blocking interventions. We tested this hypothesis in *Anopheles coluzzii* mosquitoes infected with human and rodent parasites *Plasmodium falciparum* and *Plasmodium berghei*, respectively. We find that oocyst growth rates decrease at late developmental stages as infection intensities increase; an effect exacerbated at very high infection intensities. Oocyst growth and differentiation can be restored by supplemental bloodmeals even at high infection intensities. We show that high infection intensities as well as starvation conditions reduce RNA Polymerase III activity in oocysts unless supplemental bloodmeals are provided. Our data suggest that oocysts respond to crowding and nutritional stress by employing a dormancy-like strategy and urge development of alternative methods to assess the efficacy of transmission blocking interventions.

## Main text

The lifecycle of the malaria parasite *Plasmodium* inside a mosquito vector begins with gametocytes ingested by a female mosquito during a bloodmeal. In the mosquito midgut lumen, gametes fuse to form zygotes that develop into motile ookinetes. Ookinetes traverse the midgut-epithelium and transform into oocysts. Multiple rounds of endomitotic replications inside an oocyst result in production of thousands of sporozoites, a process known as sporogony. Upon oocyst rupture, sporozoites migrate to the salivary gland for transmission to humans via a mosquito bite.

Current malaria control efforts focus on development of transmission blocking interventions including transmission blocking vaccines and drugs as well as genetically engineered mosquitoes expressing antimalarial effectors^1,2^. Experimental mosquito infections with *Plasmodium* parasites, and microscopic enumeration of oocysts is the gold standard for evaluation of transmission reduction, whereby mosquitoes are offered an infectious bloodmeal and maintained on sugar solution until oocysts are counted^3^. Infections often lead to high oocyst loads and consequently massive proliferation of sporozoites, causing depletion of nutrients in the vector^4–7^. For example, ultra-high intensity infections of *Anopheles stephensi* mosquitoes with the rodent parasite *P. berghei* leads to reduced number of sporozoites reaching the salivary glands and high mosquito mortality compared with lower infection intensities^8^. High parasite infections have been linked to marked reduction in free amino acids in the mosquito^6^, which together with lipids are chiefly acquired from bloodmeals and are essential for most physiological processes and reproduction^9^. Free amino acids are also critical for parasite development; e.g. Isoleucine (Ile), a major component of *P. falciparum* proteins, is exclusively provided by the mosquito through digestion of blood serum^10^.

Under nutritionally limiting conditions, asexual *P. falciparum* is shown to enter a dormancy-like state, restricting protein synthesis to halt cell proliferation and inhibiting catabolism to maintain viability^11,12^. Since *Plasmodium* parasites lack the target of rapamycin (TOR), the key regulator of the canonical nutrient-sensing pathway in most eukaryotes, they use the mitochondrial association factor 1 (MAF1) to regulate this metabolic switch^13^. MAF1 acts as a repressor of RNA polymerase III (Pol III) that controls expression of highly abundant non-coding RNAs and is thus essential for cell maintenance, growth and proliferation^14,15^. Under nutritional stress, *Plasmodium* MAF1 binds to Pol III switching off tRNA transcription. When conditions become favorable, MAF1 is phosphorylated and reverse translocated to the cytoplasm, freeing Pol III^13^. MAF1 mutant *P. falciparum* exhibits increased tyrosine tRNA (*tRNA*^*Tyr*^) expression and fails to recover from dormancy after long periods of incubation in Ile-deficient medium.

To date, mostly indirect evidence exists for a *Plasmodium* oocyst strategy to cope with nutritional stress. *P. berghei* oocyst size and sporozoite output are markedly reduced in *Anopheles gambiae* silenced for Lipophorin (Lp), the main lipid transporter of insects^4,16^, and as mentioned previously very high *P. berghei* infections result in reduced number of sporozoites in *A. stephensi* salivary glands^8^. These reports and unpublished observations we and others have made over the years led us to hypothesize that experimental infections may cause a nutritional stress to the developing oocyst, affecting sporogonic development and the measurable infection outcome. Here we investigated and corroborated this hypothesis using *P. berghei* and *P. falciparum* infections of *A. coluzzii* (also known as M-form *A. gambiae*). Our results stress the need for improved assays in studies of malaria transmission, especially those assessing the efficacy of transmission blocking interventions.

### Oocyst intensity affects *Plasmodium* DNA content

We optimized a quantitative PCR (qPCR) assay that detects *Plasmodium* mitochondrial *cytochrome-b* (*Cyt-b*) gene in DNA extracted from as few as 50 merozoites spiked into *A. coluzzii* midgut homogenates (**Fig. S1**). The qPCR amplification-curves were distinct from background noise in uninfected midguts (**Fig. S2**). Using this assay, we found that *PbCyt-b* abundance (used as proxy for oocyst growth) in the midgut of mosquitoes infected 7 days earlier with *P. berghei* was positively and significantly (P<0.0001) correlated with oocyst intensity recorded by microscopic enumeration prior to midgut homogenization (**Fig. 1a**). The results also showed that infection intensity does not influence *PbCyt-b* DNA abundance per capita oocyst (PCO; **Fig. 1b, c**). This pattern changes at later stages of development, as no correlation was detected between the *PbCyt-b* abundance and oocyst intensity at 14 days post infection (dpi; **Fig. 1d**). Interestingly, the oocyst intensity was negatively correlated (P=0.001) with *PbCyt-b* abundance PCO (**Fig. 1e**). When midguts were grouped into two categories based on oocyst intensity, *PbCyt-b* abundance PCO was markedly lower (P=0.007) in those harboring <15 oocysts than those with ≥15 oocysts (**Fig. 1f**).

**Fig 1.**
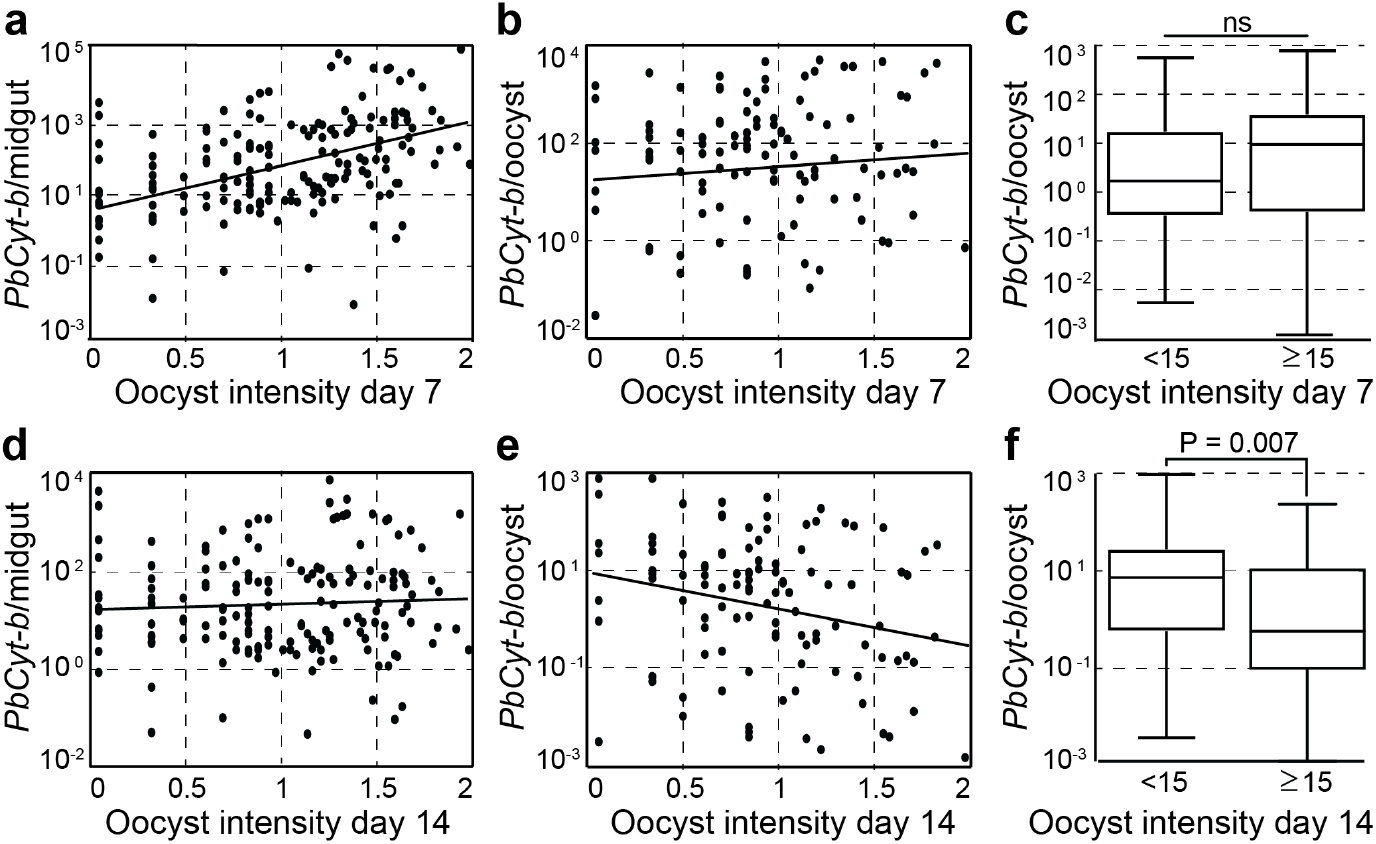
*P. berghei Cyt-b* DNA abundance is oocyst density dependent. **a**, Correlation between oocyst density and abundance of *PbCyt-b* mitochondrial DNA 7 dpi (r = 0.24; P < 0.0001). **b**, Correlation between oocyst density and *PbCyt-b* DNA abundance PCO 7 dpi. **c**, *PbCyt-b* DNA abundance PCO in low and high oocyst density midguts 7 dpi. **d**, Relationship between oocyst intensity and PbCyt-b DNA abundance 14 dpi. **e**, Correlation between oocyst density and *PbCyt-b* DNA abundance PCO 14 dpi (r = −0.25; P = 0.001). **f**, *PbCyt-b* DNA abundance PCO in low and high oocyst density midguts 14 dpi. In scatterplots, each dot represents a single midgut and lines show linear regression fit. Horizontal lines in boxplots indicate median. Data were generated from at least two independent experiments.

The impact of oocyst intensity on the abundance of parasite DNA PCO was further examined in mosquitoes where the immune factor *LRIM1* was silenced with dsRNA injection 3 days prior to *P. berghei* infection. These mosquitoes harbored over 10 times more oocysts than control *dsLacZ* injected mosquitoes (**Fig. S3**). At 14 dpi, *LRIM1*-silenced mosquitoes yielded oocysts of significantly reduced (P<0.0001) diameter compared to controls (**Fig. 2a**). A significantly decreased abundance of *PbCyt-b* PCO in *LRIM1*-silenced mosquitoes compared to controls was observed both pre-sporulation (7 dpi; P<0.0001) and sporulation (14 dpi; P<0.003) stages (**Fig. 2b, c**). These data suggested that high infection intensity negatively affects oocyst growth. Therefore, with standard post-infection mosquito husbandry, where mosquitoes obtain a single bloodmeal and are then maintained on sugar throughout the course of infection, nutrient starvation may have detrimental impacts on the developing oocyst.

**Fig 2.**
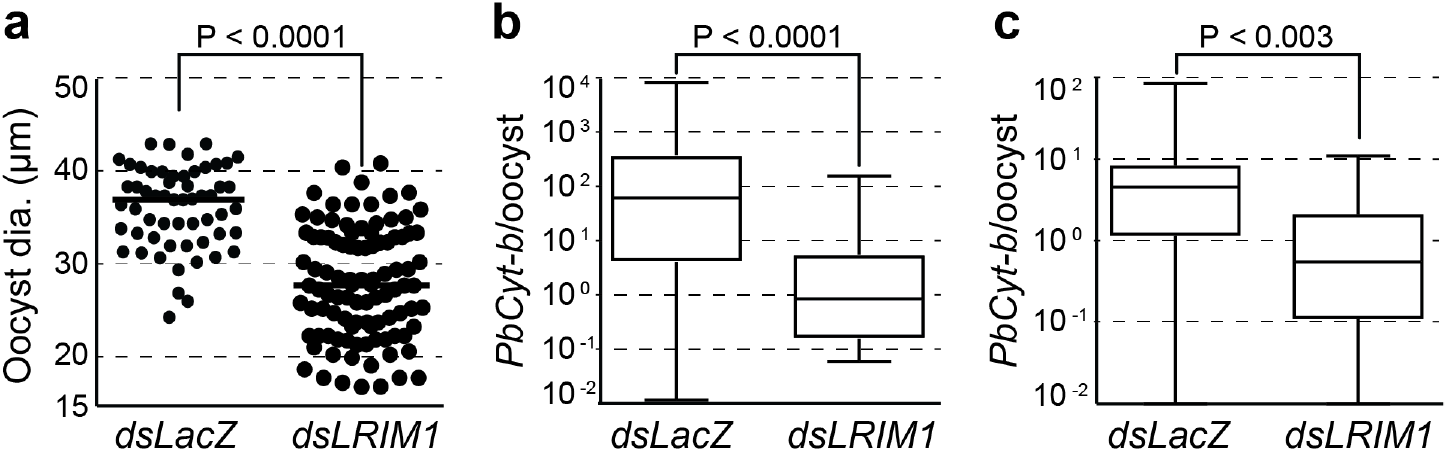
Effect of crowding on *P. berghei* oocyst growth. **a**, Oocyst diameter (dia.) in *LRIM1* and *LacZ* dsRNA injected mosquitoes. Data were generated from two independent biological replicates. **b-c**, *PbCyt-b* DNA abundance in *LRIM1* and *LacZ* dsRNA injected mosquitoes 7 (b) and 14 (c) dpi. Data were generated from three independent replicates. Horizontal lines in dot plots and boxplots indicate median.

### Supplemental bloodmeals accelerate oocyst growth

To assess the impact of post-infection mosquito blood feeding on oocyst growth, parasite DNA abundance was compared between *P. berghei* oocysts in mosquitoes maintained under standard husbandry regime and mosquitoes provided a supplemental, uninfected bloodmeal 7 dpi. The results revealed that the supplemental bloodmeal significantly increased (P=0.029) the oocyst size compared to control (**Fig. 3a, b**). In these mosquitoes, oocysts contained on average significantly more DNA (P=0.003) than oocysts in control mosquitoes (**Fig. 3c**). The supplemental bloodmeal significantly increased (P<0.0001) the correlation between *P. berghei* oocyst intensity and DNA abundance per midgut (**Fig. 3d**). It also abolished the negative correlation between oocyst intensity and abundance of parasite DNA PCO (**Fig. 3d**), which was observed in mosquitoes under standard husbandry regime (see **Fig. 1e**), revealing that oocyst growth ensues when mosquitoes receive a supplemental bloodmeal even when infection intensities are high.

**Fig 3.**
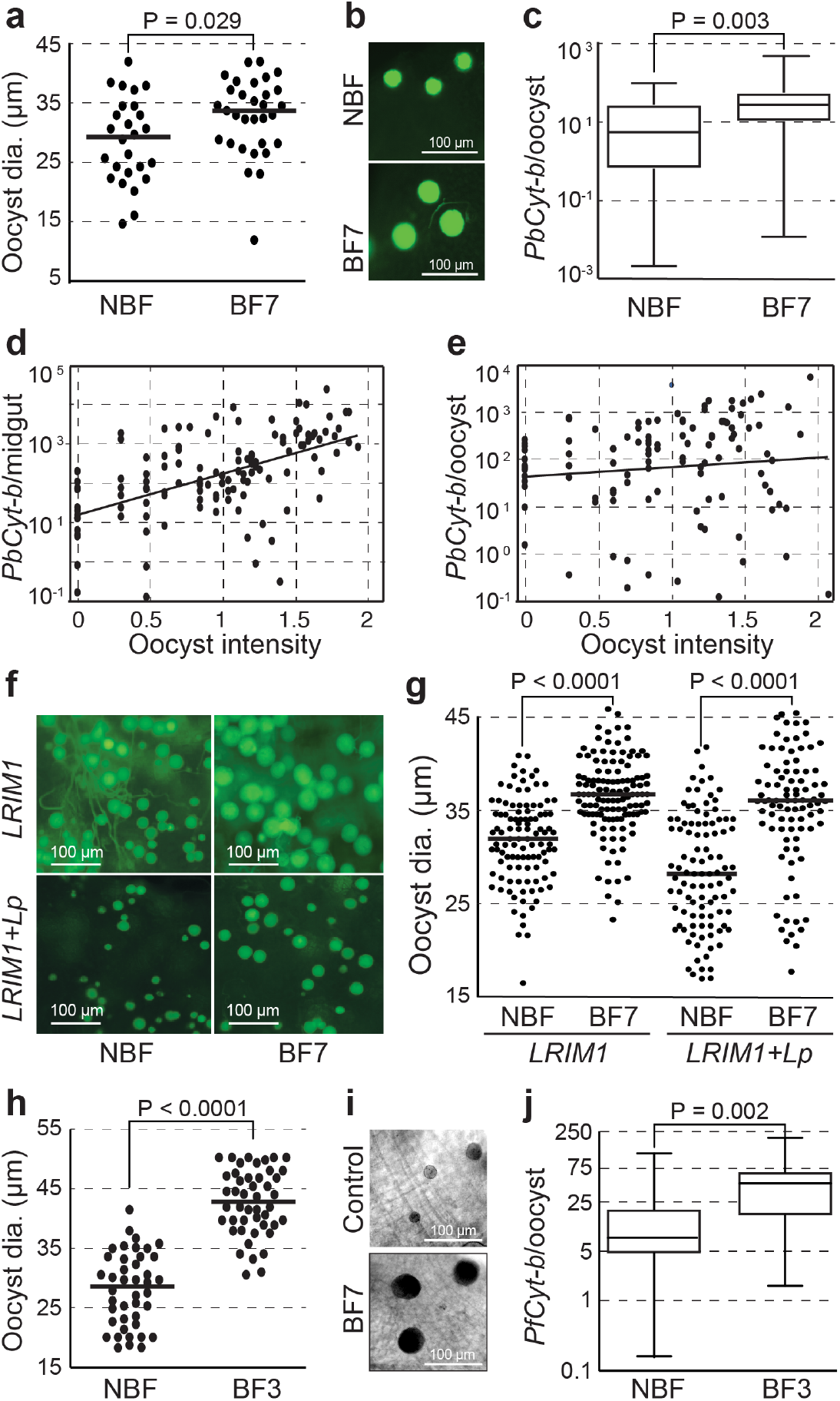
Effect of supplemental bloodmeals on oocyst growth. **a**, *P. berghei* oocyst diameter (dia.) in mosquitoes provided no supplemental bloodmeal after the time of infection (non-blood-fed; NBF) or an additional supplemental bloodmeal 7 dpi (BF7) measured 14 dpi. **b**, Representative microscopic pictures of *P. berghei* oocysts in NBF and BF7 mosquitoes taken 14 dpi. **c**, *PbCyt-b* DNA abundance in NBF and BF7 mosquitoes 14 dpi. **d**, Correlation between *P. berghei* oocyst density and PbCyt-b DNA abundance in the midgut of NBF and BF7 mosquitoes (r = 0.35, P<0.0001). **e**, Correlation between *P. berghei* oocyst density and *PbCyt-b* DNA abundance PCO. **f-g**, Representative microscopic pictures (**f**) and diameter (**g**) of *P. berghei* oocysts on midguts of mosquitoes silenced for *LRIM1* and *LRIM1*+*Lp* and maintained under a standard husbandry regime (NBF) or provided a supplemental bloodmeal 7 dpi (BF7). For oocyst diameter, oocysts were randomly selected from 20 midguts from two independent infection experiments. **h**, *P. falciparum* oocyst diameter (dia.) in mosquitoes provided no supplemental bloodmeal after the time of infection (NBF) or an additional supplemental bloodmeal 3 dpi (BF3) measured 9 dpi. **i**, Representative microscopic pictures of *P. falciparum* oocysts in NBF and BF3. **j**, *PfCyt-b* DNA abundance in NBF and BF3 mosquitoes. Horizontal lines in dot plots and boxplots indicate median. Data were generated from at least two independent experiments.

We further investigated the effect of mosquito starvation on oocyst growth by augmenting oocyst numbers through silencing *LRIM1* either alone or in combination with *Lp*, thus creating extreme crowding and starvation conditions^4^. The results revealed a remarkable variation of *P. berghei* oocyst size in midguts of both *LRIM1* and *LRIM1*+*Lp* silenced mosquitoes maintained under a standard husbandry regime, particularly in the latter mosquito cohort (**Fig. 3f** **left panels**). This phenotype was reversed when mosquitoes were provided a supplemental bloodmeal 7 dpi, as shown by a significant increase of median oocyst size in both *LRIM1* (P<0.0001) and *LRIM1*+*Lp* (P<0.0001) silenced mosquitoes (**Fig. 3f** **right panels** and **Fig. 4b**).

**Fig 4.**
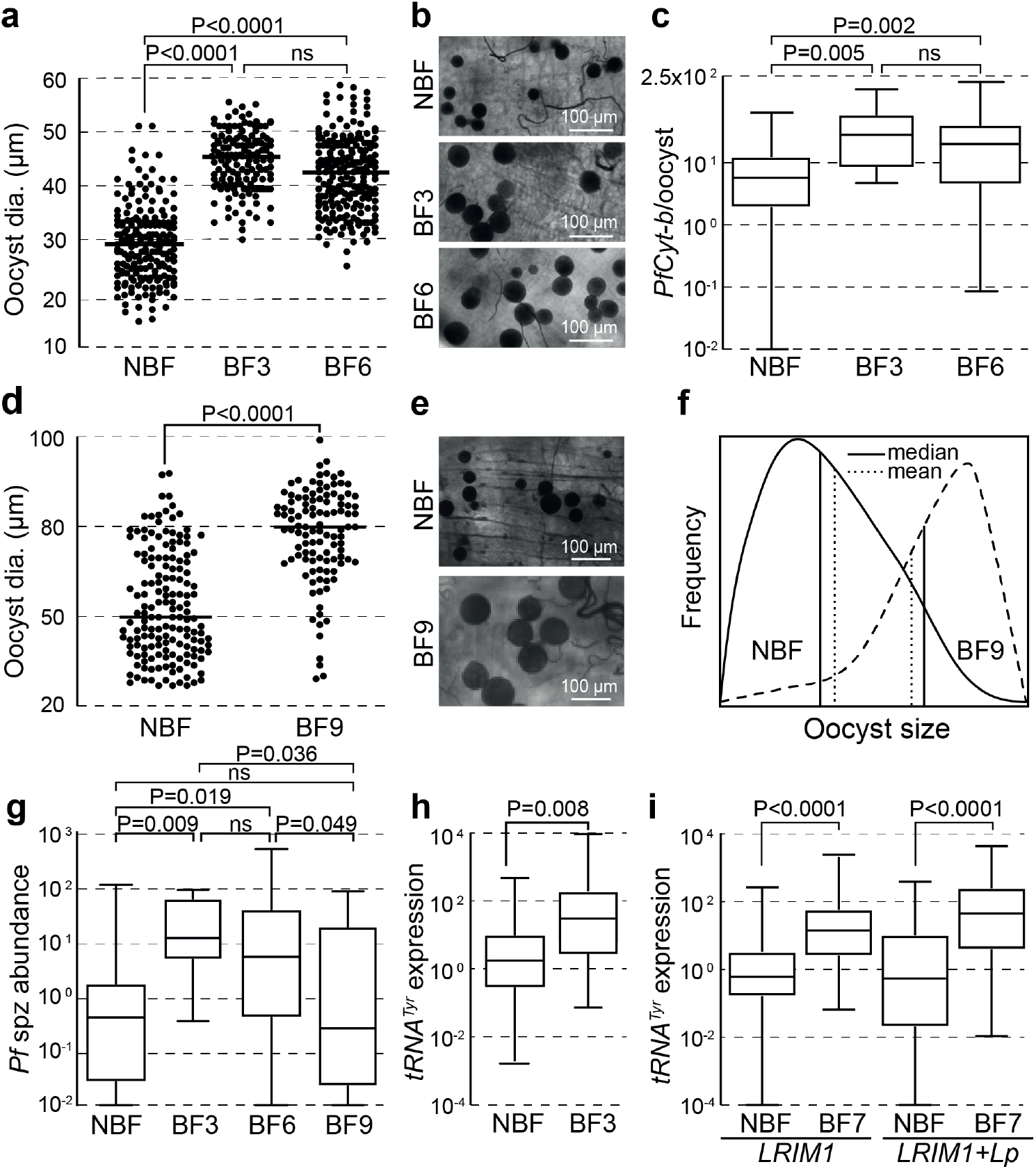
Effect of supplemental bloodmeal deprivation on oocyst development. **a**, *P. falciparum* oocyst diameter (dia.) in mosquitoes provided no supplemental bloodmeal after the time of infection (non-blood-fed; NBF) or additional supplemental bloodmeals at 3 (BF3) or 6 (BF6) dpi measured 9 dpi. **b**, Representative microscopic pictures of *P. falciparum* oocysts in NBF, BF3 and BF6 mosquitoes taken 9 dpi. **c**, *PfCyt-b* DNA abundance in NBF, BF3 and BF6 mosquitoes 9 dpi. **d**, *P. falciparum* oocyst diameter (dia.) in mosquitoes provided no supplemental bloodmeal after the time of infection (NBF) or additional supplemental bloodmeals 9 dpi (BF9) measured 13 dpi. **e**, Representative microscopic pictures of oocysts in NBF and BF9 mosquitoes taken 13 dpi. **f**, Oocyst size distribution fitting the Poisson distribution model in NBF and BF9 mosquitoes 13 dpi. Solid vertical lines show median and dotted vertical lines show mean. **g**, Sporozoite load in NBF, BF3, BF6 and BF9 mosquitoes at 14 dpi. **h**, Relative expression levels of *P. falciparum pre-tRNA*^*Tyr*^ in NBF and BF3 mosquitoes measured 7 dpi. **i**, Relative expression levels of *P. berghei pre-tRNA*^*Tyr*^ in *LRIM1* and *LRIM1*+*Lp* silenced mosquitoes in NBF and BF7 mosquitoes at 14 dpi. Horizontal lines in dot plots and boxplots indicate median. Data were generated from at least two independent biological replicates.

Analogous and indeed more pronounced differences were recorded for *P. falciparum* oocysts in mosquitoes that received a supplemental bloodmeal at day-3 pi compared to control mosquitoes that received only a single, infected bloodmeal (**Fig. 3h-j**). The supplemental bloodmeal was provided at day-3 pi due to the faster bloodmeal digestion and oocyst development in these mosquitoes which are maintained at 27°C instead of 21°C, while dissection was performed at day-9 pi.

### Oocysts enter a dormancy-like state under starvation conditions

Female mosquitoes naturally lay their eggs before taking another bloodmeal to replenish energy reserves and embark on a new gonotrophic cycle. Follicular resorption that is mediated by juvenile hormone is employed to increase energy resources in case of nutrient limitation or when oviposition is not possible^17^, as is the case of most experimental infection protocols. This process is presumed to also provide nutrient supplies to the developing oocyst^8,18^. We reproduced the natural infection conditions by allowing mosquitoes infected with *P. falciparum* to lay their eggs at day 2 pi, before they were provided supplemental bloodmeals at either day 3 or day 6 pi. The results showed a dramatic difference reflected by significantly decreased oocyst sizes recorded at day 9 pi in mosquitoes deprived of either of supplemental bloodmeals (both P<0.0001; **Fig. 4a, b**). This finding was consistent with lower parasite DNA abundance PCO in bloodmeal-deprived mosquitoes compared to those provided a supplemental bloodmeal at 3 (P<0.005) or 6 (P<0.002) dpi (**Fig. 4c**). No significant differences in oocyst size or parasite DNA abundance PCO were recorded between mosquitoes provided supplemental bloodmeals at 3 or 6 dpi, suggesting that oocyst growth can quickly recover after a bloodmeal.

Oocysts retained their propensity to grow even when a supplemental bloodmeal was provided as late as 9 dpi (P<0.0001; **Fig. 4d, e**). In this experiment, midgut dissections and oocyst observations were carried out 13 dpi. Interestingly, the distribution of oocyst sizes in mosquitoes deprived of supplemental bloodmeal was positive-skewed (**Fig. 4f**), and several oocysts in these mosquitoes were as big as the biggest oocysts in mosquitoes provided a supplemental bloodmeal 9 dpi. This implies that, under limiting nutrient conditions, the oocyst population employs an effective resource management strategy whereby some oocysts are let to grow whereas most remain dormant. When faced with resource stress, *Plasmodium* is indeed thought to induce growth restriction mediated by extracellular vesicles, limiting growth to some parasite cells and ensuring transmission^19^. Furthermore, the distribution of oocyst sizes in mosquitoes receiving a supplemental bloodmeal 9 dpi was negative-skewed, and several oocysts were as small as the smallest oocysts in mosquitoes deprived of supplemental bloodmeals, suggesting that not all oocysts effectively exit dormancy following a long starvation period.

We investigated the abundance of *P. falciparum* sporozoites in salivary glands 14 dpi by quantifying *PfCyt-b* in the mosquito thorax and head regions (**Fig. 4g**). The results revealed that deprivation of supplemental bloodmeals combined with oviposition at 2 dpi resulted in significantly lower levels of sporozoites compared to mosquitoes provided supplemental bloodmeals at 3 (P=0.009) or 6 (P=0.019) dpi. Nevertheless, most of these mosquitoes (70%) did harbor detectable sporozoite loads in their salivary glands, like those provided supplemental bloodmeals (81% at 3 dpi and 86% at 6 dpi), indicating that some oocysts, presumably the fully-grown ones, were fully differentiated and produced sporozoites. Despite having much larger oocysts, mosquitoes provided a supplemental bloodmeal 9 dpi, showed equal sporozoite loads compared to those deprived of supplemental blood meals, suggesting that more time was needed for full oocyst differentiation.

The decelerated oocyst growth in nutrient limiting conditions suggested that oocysts might employ a dormancy-like strategy to overcome starvation. To examine this, parasite *pre-tRNA*^*Tyr*^ abundance was quantified as a proxy for Pol III activity in oocysts. Measuring immature instead of mature *tRNA*^*Tyr*^ served to overcome the long half-life of mature tRNA that might affect real-time quantification of gene expression^13^. *Pre-tRNA*^*Tyr*^ expression was calibrated to parasite DNA abundance PCO. The results revealed that the levels of *P. falciparum pre-tRNA*^*Tyr*^ expression levels in oocysts 7 dpi were significantly lower (P=0.008) in mosquitoes that received a single, infectious bloodmeal compared to those that additionally received a supplemental bloodmeal 3 dpi (**Fig. 4h**). Similarly, *P. berghei tRNA*^*Tyr*^ expression in *LRIM1* (P<0.0001) and *LRIM1*+*Lp* silenced (P<0.0001) conditions measured in oocysts 14 dpi were significantly lower in mosquitoes under standard husbandry regimes compared to those that received a supplemental bloodmeal 7 dpi (**Fig. 4i**). These data together suggest that oocysts enter a dormancy-like state in the mosquito vector when faced with nutrient depletion caused by bloodmeal deprivation, high infection intensities or disruption of nutrient shuttling.

## Discussion

*Plasmodium* spends a large part of its life as an oocyst within the *Anopheles* mosquito, where it faces a variety of challenges that can adversely affect its growth, differentiation and transmission potential^20^. A key challenge is the high nutritional demand to produce thousands of sporozoites. We demonstrate that under starvation conditions, when an oocyst-harboring mosquito is deprived of supplemental bloodmeals, infection intensity plays a major role in oocyst growth, as measured by both DNA content and oocyst size. Mosquitoes harboring many oocysts, a condition that can occur naturally in field infections and invariably in experimental infections and which can be further elevated by inactivating the mosquito immune response, exhibit substantial variability in oocyst growth. In such conditions of crowding and nutritional stress, expression of parasite *pre-tRNA*^*Tyr*^ is significantly reduced, indicating that the oocyst enters a dormancy-like state.

Our findings bear important implications not only for understanding the mosquito-parasite interactions underpinning malaria transmission but also in evaluating the efficacy of transmission blocking interventions. The current gold standard in malaria transmission studies is microscopic enumeration of oocysts present in the mosquito midgut. It has been previously shown that the number of oocysts correlates well with the number of gametocytes ingested by the mosquito and can therefore be used as a measure of the efficacy of interventions acting on gametocytes, gametes and/or ookinetes^21–23^. However, this measurement does not consider any knock-on or additional impacts of the intervention on the oocyst. Although this additional impact is unlikely to reverse any detrimental effect on previous developmental stages, our data suggest that reduction in oocyst numbers may translate to a lesser or no impact on the sporozoite load, especially when the effect is small and oocyst competition for nutrients is reduced. Thus, a reduction in oocyst intensity alone is an insufficient measurement and may not accurately reflect the efficacy of a transmission blocking intervention.

At high intra-population densities, endoparasites are known to evoke competition for nutrients when these are of limited supply^24^. This phenomenon is ubiquitous in nature and often leads to negative density-dependent processes with significant effects on parasite fitness such as proliferation and ability to avoid host immune defenses^25^. The density-dependent effect on *Plasmodium* sporogony reported here is not related to mosquito immune responses known to reduce the number of oocysts but to the rate of oocyst growth through endoreplication. The effect is indeed exacerbated upon silencing the mosquito complement-like system due to the significant increase of the number of ookinetes that successfully develop to oocysts.

The negative density-dependent effect on the oocyst is exacerbated in standard laboratory infections where infected mosquitoes are often not provided supplemental bloodmeals, leading to acute depletion of nutrient reserves^4–7,26^. In nature, a female mosquito acquires a bloodmeal every 2-3 days^27,28^, which helps it meet the high energy and nutrient demand for flight and reproduction. These additional bloodmeals are also thought to be important for the proliferating oocyst^18^. *Plasmodium* is auxotrophic for most amino acids; of these, free Isoleucine exclusively derives from bloodmeal digestion, as neither the parasite nor the mosquito can synthesize it *de novo*^*10,29,30*^.

Bloodmeal-derived fatty acids are also critical for mosquito fitness and reproduction, chiefly derived from hemoglobin metabolism in the gut cells and distributed across mosquito tissues by Lp. Silencing the *Lp* gene abolishes egg development and compromises growth of both *P. berghei* and *P. falciparum* oocysts^4,16^; albeit studies showing that mature late-stage *P. falciparum* oocysts synthesize lipids de novo via the type two fatty acid biosynthesis pathway (FAS-II), in contrast to *P. berghei* and asexual stages that scavenge fatty acids exclusively from the host^31^. We show that silencing *Lp* together with the complement-like immune pathway exacerbates the negative effects of crowding on oocyst growth but cannot further reduce *pre-tRNA*^*Tyr*^ expression, indicating that the dormancy-like state is likely to be regulated by amino acid sensing^12^.

Better understanding of the oocyst response to crowding and nutritional stress could assist development of novel interventions to limit malaria transmission. For example, a temporally tuned targeting of pathways involved in mosquito amino acid or lipid metabolism using genetic engineering could prolong completion of the parasite extrinsic incubation period (EIP), delaying oocyst growth and differentiation and sporozoite release. In nature, only a small proportion of mosquitoes can survive through the *P. falciparum* EIP^32,33^; hence, even a small delay in oocyst growth and differentiation could have a drastic negative impact on malaria transmission.

Importantly, our findings highlight a major drawback of most standard laboratory infection protocols that can lead to erroneous conclusions when evaluating the transmission-blocking efficacy of an intervention. First, an intervention that inhibits or delays oocyst growth and/or differentiation without or by weakly affecting oocyst counts may be rejected owing to a decision merely based on reduction of oocyst numbers. This pertains to interventions targeting both mosquito or parasite molecules and pathways, e.g. mosquito *Lp* as described earlier^16,34^ and parasite *misfit* gene of which the knockout produces oocysts arrested at early mitotic stages but continuing to grow in size^35^. Second, inducing a dormancy-like state, a universally resilient life stage, may make oocysts resistant to transmission blocking interventions. Third, the current standard protocol may lead to false positive conclusions in favor of an intervention reducing oocyst numbers whilst it has no real impact on malaria transmission owing to the density-dependent effects described here. We conclude that provision of supplemental bloodmeals to mitigate any parasite density-dependent effects on oocyst growth and development and assessment of the actual parasite load in the oocyst that escape the intervention through a molecular diagnostic method are profoundly important.

## Methods

### Mosquito infections

*A. coluzzii* of the N’gousso strain were reared following standard protocols and infected with *P. berghei ANKA507m6cl1* that constitutively expresses GFP^36^ by direct feeding on infected mice or with *P. falciparum* gametocytes from a laboratory culture (NF54 strain) using standard membrane feeding. Engorged mosquitoes were provided 10% sucrose and maintained at 21°C for *P. berghei* infections and 27°C for *P. falciparum* infections until midgut dissection. Supplemental bloodmeals on human blood was provided via membrane feeding.

### Midgut preparations and oocyst enumeration

Gut dissections were performed in ice-chilled PBS. For *P. berghei* infection, guts were mounted on glass slides and oocysts were counted with fluorescence microscopy. For *P. falciparum* infection, guts were stained with 0.5% Mercurochrome for 10 min on ice and washed twice in ice chilled PBS for 30 min, and oocysts were enumerated using light microscopy. After oocyst enumeration, the foregut and hindgut were removed, and midguts were individually transferred to 1.5 ml screw-top tubes with 350 μl RLT Plus buffer from Qiagen AllPrep DNA/RNA Mini Kit and kept at −20°C until DNA and RNA extraction.

### Genomic DNA and RNA extraction

Midguts were homogenized using PRECELLYS® 24 (Bertin Technologies) before simultaneous extraction of genomic DNA and total RNA using the AllPrep DNA/RNA Mini Kit. DNA and RNA concentrations were determined using Nanodrop (Thermo Scientific), and samples were kept at −20°C until cDNA synthesis and qPCR analysis. To measure the abundance of *P. falciparum* sporozoites in salivary glands, mosquitoes were killed with 75% ethanol and placed on dry ice and their abdomens were removed. Genomic DNA was extracted from the remaining thorax and head regions of each mosquito using Qiagen DNeasy 96 Blood and Tissue kit.

### Quantitative PCR analysis

SYBR Green based qPCR was used to quantify *Plasmodium Cyt-b* mitochondrial DNA. Assay sensitivity and specificity were determined with serial dilutions of *P. falciparum* merozoites from a synchronized asexual culture after heat inactivation, quantified and collected using a BD FACSAria III cell sorter. QPCR reactions were performed in 20 μl volumes according to the manufacturer’s instructions and previously described primers for *P. berghei*^37^ and *P. falciparum*^38^. *Pre-tRNA*^*Tyr*^ expression levels were determined with the Qiagen Quantinova SYBR Green PCR kit using the stem-loop qPCR technique and qPCR primers described previously^13^.

DNA and RNA abundances were measured using the standard curve-based qPCR method. Curves for each target gene and the reference gene (*A. gambiae* S7 ribosomal protein gene^39^) were constructed after serial dilution of nucleic acid templates. Ct-values were standardized using their respective standard curves, and the target gene Ct value was normalized to that of the reference gene. *Pre-tRNA*^*Tyr*^ PCO was standardized to *Cyt-b* abundance.

### Data analysis

GraphPad Prism v.8.4.0 software package was used for data analysis and presentation. Spearman’s rank-order correlation analysis was used to measure the relationship between oocyst intensity (log_10_ transformed) and relative DNA abundance per single midgut or oocyst. All other statistical analyses were performed using the Kolmogorov-Smirnov test.

## Acknowledgements

We thank Jane Srivastava and Jessica E. Rowley for assisting with parasite cell sorting and Maria G. Inghilterra for gametocyte culturing. The work was funded by the Bill and Melinda Gates Foundation grant OPP1158151 to N.W. and G.K.C and the Wellcome Trust Investigator Award 107983/Z/15/Z to G.K.C.

## Author contributions

T.H. and G.K.C. contributed conceptualization; T.H., A.A.S., C.A.S.W. and E.K.G.M. contributed methodology; T.H. and A.A.S. conducted investigation; T.H. and G.K.C. performed formal analysis; T.H. and A.A.S. wrote original draft; T.H. and G.K.C. wrote the paper; T.H., N.W. and G.K.C. provided supervision; G.K.C. conducted project administration; N.W. and G.K.C. acquired funding.

## Supplementary Figures

**Fig S1.**
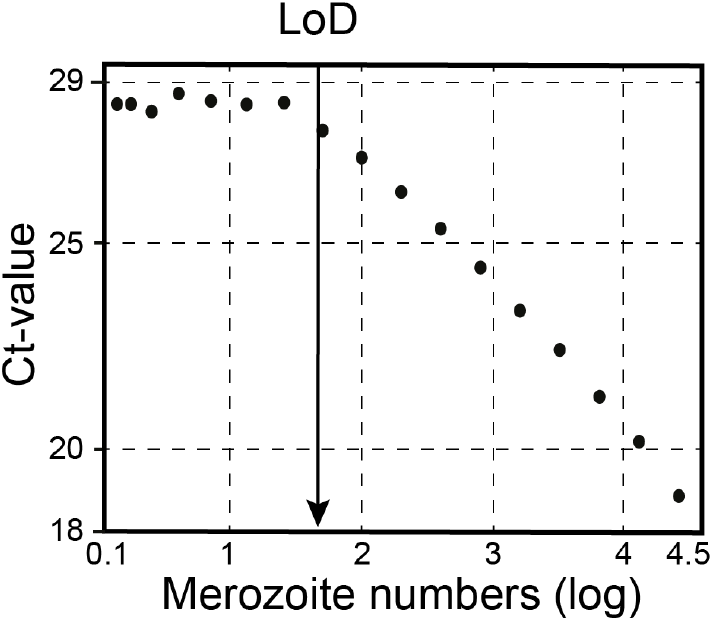
Efficiency of quantitative PCR in determining *PfCyt-b* DNA abundance. Merozoite cells from *P. falciparum* asexual cultures were purified and quantified using FACS sorter. Single midgut homogenates from primiparous *A. coluzzii* mosquitoes were spiked in merozoite serial dilutions. SYBR-based qPCR was performed following DNA extraction. The standards curve represent average of 15 independent qPCR assays. Arrow indicates the limit of assay detection (LoD). Note that each merozoite cell is thought to contain up to 20 mitochondria.

**Fig S2.**
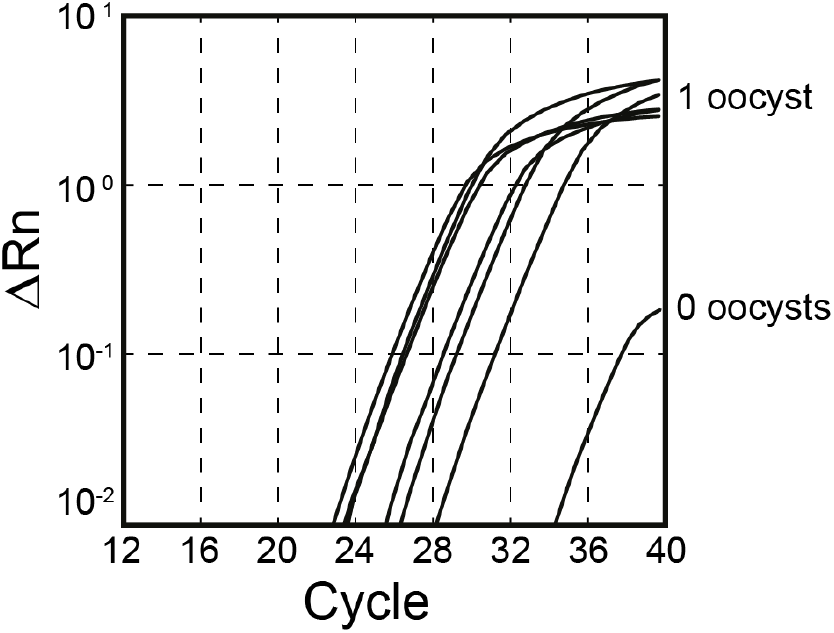
*PfCyt-b* DNA qPCR amplification curves in midguts with single or no oocysts. Midguts of *A. coluzzii* mosquitoes fed on *P. falciparum* gametocytemic blood were dissected at 7 dpi and the number of oocysts were determined. DNA was extracted from 6 midguts showing mono-oocyst infections and 1 with no oocysts. Abundances of *PfCyt-b* DNA in midguts containing a single oocyst in relation to mosquito *S7* gene are 3.24, 6.14, 17.86, 40.37, 0.99, 0.92 and 0.002 from left to right, respectively.

**Fig S3.**
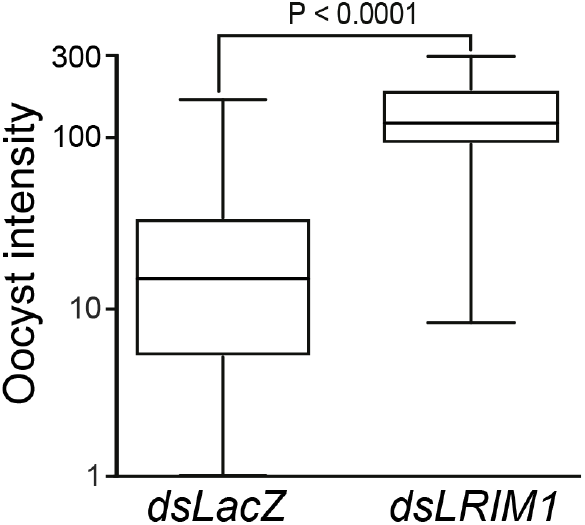
*P. berghei* oocyst intensity following silencing of *LRIM1*. Oocysts present on mosquito midguts were counted at day 14 pi. Injection of *LacZ* dsRNA was used as control. Horizontal lines in boxplots show median. Data were generated from three independent biological replicates. Kolmogorov-Smirnov test was applied to compare oocyst counts between *dsLacZ*-injected and *LRIM1* silenced mosquitoes.

